# The illusion of infinity: Acoustic black holes in wood deceive termites

**DOI:** 10.1101/2025.10.13.681988

**Authors:** Can Nerse, Shahrokh Sepehrirahnama, Joseph C.S. Lai, Theo A. Evans, Sebastian Oberst

**Affiliations:** Centre for Audio, Acoustics and Vibration, Faculty of Engineering and Information Technology, University of Technology Sydney, Australia; School of Engineering and Technology, University of New South Wales, Australia; School of Animal Biology, University of Western Australia, Australia

## Abstract

Vibrational signals produced during feeding are fundamental to termite behaviour, yet their function in regulating collective foraging remains unclear. In this study, we combine bioassays, micro-CT imaging, and elastic wave modelling to investigate how the subterranean termite *Coptotermes acinaciformis* evaluates wood through structural wave propagation. Using an axially excited Acoustic Black Hole (ABH), a tapered geometry that minimises wave reflections and effectively mimics an infinitely long food source, we show that termites preferentially attack longer wooden dowels and, remarkably, also lighter ABH-modified dowels. Micro-CT scans revealed feeding concentrated in the dowel core, coinciding with the region of maximum stress predicted by the models but where echo return was minimal. These results indicate that termites assess wood size through bite-induced echoes, analogous to echolocation in bats and dolphins, and preferentially exploit core regions of trunks and branches, thereby accounting for the tree-piping behaviour of termites. The reduction or absence of reflected waves may thus act as a cue that stimulates collective stigmergic foraging. From an applied perspective, ABH-inspired structures could form the basis of novel, chemical-free lures for termite management.

---

Mechanoreceptors, widespread across insect orders, also play a crucial role in how termites perceive and interact with their environment. Yet, unlike many other insects, termites are blind and lack specialized sensory structures such as tympanal organs for detecting airborne sound. Instead, termites use substrate-borne vibrations generated during their wood-biting activity to assess the quantity and quality of the wood (*1, 2*), to make foraging decisions (*2, 3*), to distinguish kin (*2*) and other species, including competitors (*4*) and predators (*5*), and to signal alarm (*6, 7*).

Among insects that directly feed on wood, *Coptotermes* have a distinctive foraging behavior; they evolved to nest and eat trees from within and are therefore also called *tree-piping termites*. They target the central axis of the tree, hollowing out the trunk and the branches, while leaving the sapwood and bark intact (*8*). It is well established that termites use vibrations to guide foraging decisions; however, the specific signal features underpinning this tree-piping capability remain poorly understood.

In solids such as wood and clay, vibrations propagate as elastic waves (*9*). When these waves encounter a discontinuity, such as a boundary or structural imperfection, they are reflected back to the point of origin as ‘echoes’. At resonant frequencies of the material, successive reflections interfere constructively, amplifying the signal.

While termites could, in principle, use amplitude as a cue for foraging (*1*), the eigenfrequency of wooden structures appears not to play a dominant role (*3, 10*). Instead, termites rely on specialised subgenual sensilla located in the tibiae of their legs (*11*), which make them highly sensitive to transient vibration signals (*12*). They can also determine the direction of a stimulus by detecting phase differences between their legs (*13*), and they also respond to static stress such as the weight load of a tree (*14*).

It is therefore plausible that termites biting into wood generate stress waves whose echoes they detect through amplitude and phase. This mechanism may provide the missing link in explaining how termites assess food quantity and quality using vibrational cues.

To test this hypothesis, we conducted bioassays with *Coptotermes acinaciformis* using *Pinus radiata* dowels of varying length and diameter to examine foraging preferences (Fig. 1*A*). We then modelled elastic wave propagation and echo formation using finite-element explicit time-domain simulations to elucidate potential perception mechanisms. In addition, we introduced dowels with one end shaped into an Acoustic Black Hole (ABH), a taper that delays and reduces the amplitude of echoes (*15*), to directly test whether termites perceive and use echoes when making foraging decisions.

**Figure 1:**
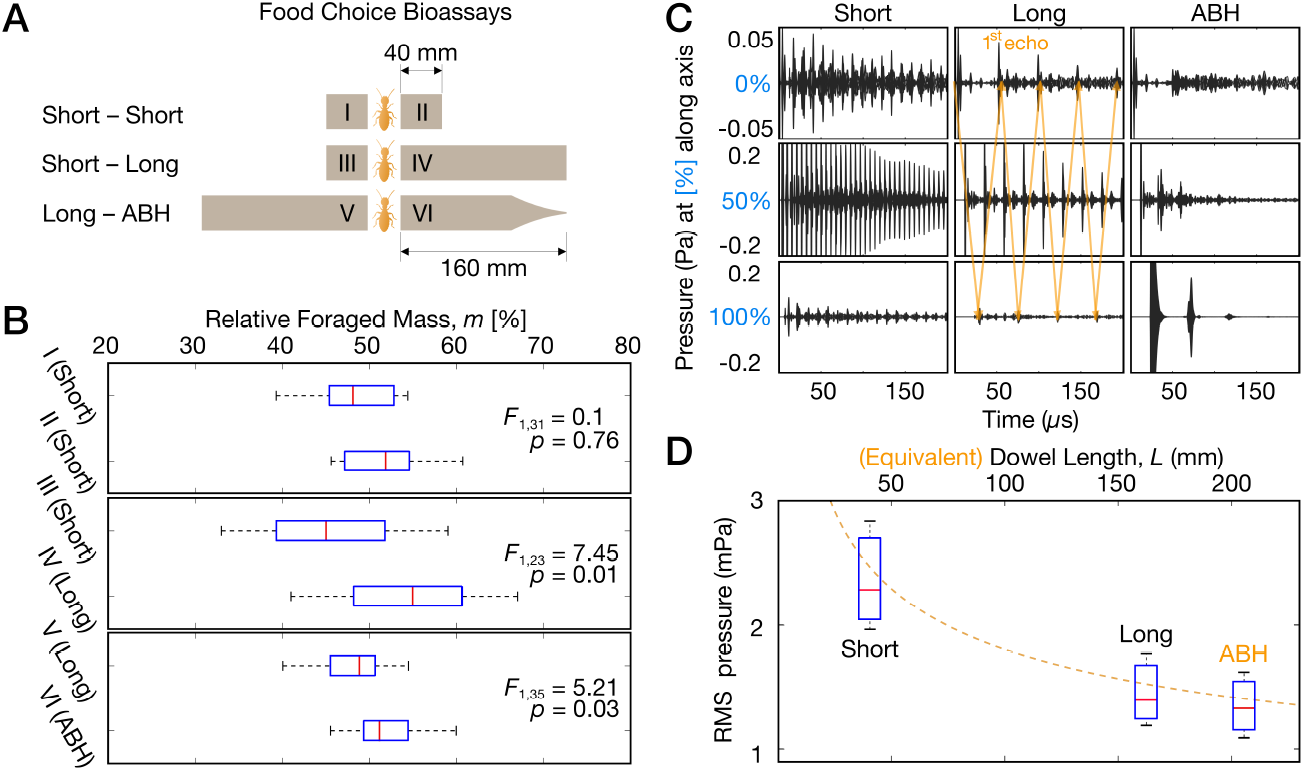
(A) Food-choice bioassay design, using *N* = 25 samples per control (Short *P. radiata* dowels) and treatment for seven colonies. Treatments consist of a long dowel and an Acoustic Black Hole (ABH) termination to investigate preference for larger food resources. (B) Statistical analysis of the percentage of the foraged mass of the *P. radiata* dowels. The *F*-values and their respective *P*-values from the one-way ANOVA tests are shown for the control of each treatment. (C) Elastic wave propagation due to a unit pressure pulse mimicking termite biting on the wood (*SI Appendix*) is depicted by pressure signals at defined positions along the axis. (D) Root mean square (RMS) pressure at the termite-feeding end, i.e., 0%, is used to evaluate whether termites sense their biting signals. Time evaluation period (Δ*t*) is selected concerning estimated termite temporal sensing threshold (0.1−0.3 ms) (*13*). The length and mean RMS pressure ratio of the short and long dowels are used to calculate the equivalent length of the ABH dowel. For Δ*t* = 0.3 ms, in a 4:1 length ratio, the pressure ratio is calculated as 0.6 (1.2:2.0 mPa), thus varying with 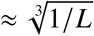 (dashed line), the ABH dowel with mean RMS pressure of 1.1 mPa is used to deceive the termites as it approximately generates the same pressure level as a piece that is 205 mm long.

## 1 Results

### 1.1 Termites Infer Equivalent Length from Amplitude of Stress Waves

In the bioassays, using the relative dry weight of consumed wood, termites showed no preference with two paired short dowels (I + II — both 40 mm, Fig. 1*B*). Further, they preferred long (IV = 160 mm) over short dowels (III = 40 mm). However, termites preferred the ABH design over long dowels, even though the ABH dowels had about 14% lower mass (due to milling of the dowel to shape the ABH end).

In the bioassays, using the relative dry weight of consumed wood, termites showed no preference with two paired short dowels (I + II — both 40 mm, Fig. 1*B*), and they preferred long (IV = 160 mm) over short dowels (III = 40 mm). However, termites preferred the ABH design over long dowels, even though the ABH dowels had about 14% lower mass (due to milling of the dowel to shape the ABH end).

### 1.2 Stress Waveform Gives Insights on Foraging Path

We examined termite feeding in the dowels with *μ*CT scans. In all cases, termites excavated a cone shape, so their feeding was concentrated at an apex in the center of the dowel (Fig. 2*A*). This pattern suggested termites may be following a signal at the center of the dowel, which led to our next inquiry. We modeled the vibration signals, created by termite chewing and echoed by the distal end of the dowels (Fig. 2*B*).

**Figure 2:**
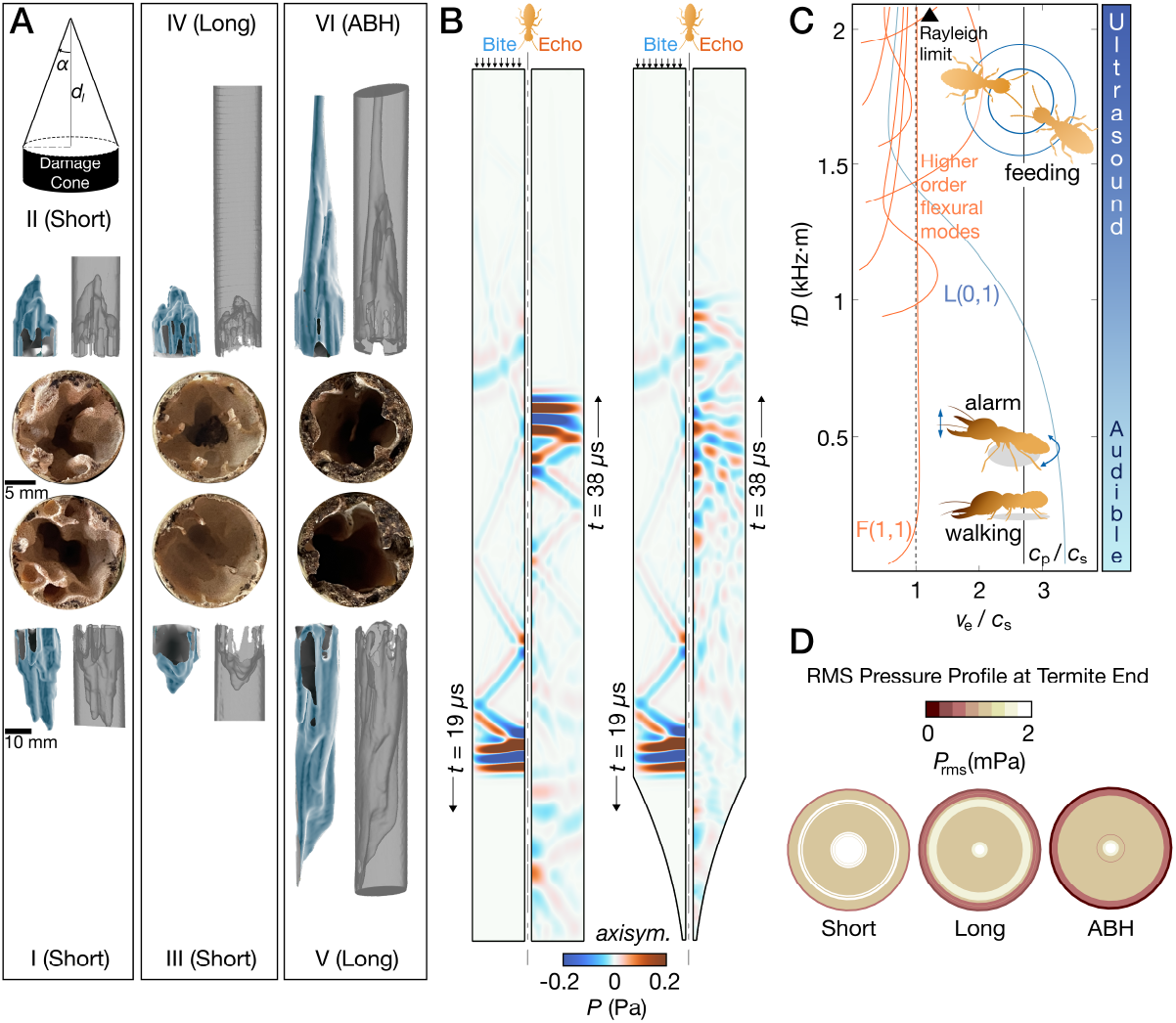
(A) *μ*CT scans of a subset of damaged dowels and the cone-shaped damage pattern. The internal damage pattern in all cases resembles a cone, which can be quantified by the depth *d*_*l*_ and the cone angle *α* of the damage frontier. (B) Elastic wave propagation for a uniform and ABH dowel is shown at time *t*_1_ = 19 *μ*s, and *t*_2_ = 38 *μ*s by pressure distribution in an axisymmetric dowel configuration. The direction of the arrow corresponds to the wavefront before (down) or after (up) the first reflection from the tip of the dowels. While the wavefront for the uniform dowel is constant, the ABH dowel shows a stochastic pattern due to thickness variation. With each reflection, |*R*| *<* 1, flexural and longitudinal wave families slow down along the tapered end. This results in the accumulation of strain energy at the tip of the ABH dowel, which can be explained by the asymmetric pressure time history at the ends of the dowel, as shown in Fig. 1*C*. Energy velocity dispersion curves for a perfectly elastic *P. radiata* dowel with *D* = 19 mm are shown in (C). Commonly observed termite behaviors are denoted in terms of the frequency range and wave families associated with their acoustic emission. (D) At a typical frequency range for feeding-induced acoustic emission (*16*), numerically obtained stress profiles at the dowel end at the feeding site predict higher pressure levels at the inner core of the dowels, especially for the ABH dowels along *r* = 0 for a pulse input signal. These results correlate with the damage cone in the dowel samples used in bioassays, shown in (A), where termite tunneling activity is greater at the regions associated with higher dynamic stress.

For the long dowel, more vibration energy propagated longitudinally, resulting in weaker shear waves and a more uniform pressure distribution across the dowel’s end (Fig. 2*B*). In contrast, the ABH dowel exhibited the lowest overall pressure level (Fig. 1*C* and *D*), with a non-uniform distribution (Movie S1). This effect is primarily attributed to the ABH profile, which enhanced shear wave generation and trapped vibration energy at the ABH section as a lateral standing wave (Fig. 2*C*), thus reducing the maximum pressure signal amplitude of the dominant wavefront (string of echoes) and contributing to increased energy dissipation. The strongest pressure signal occurs at the inner core of dowels (near *r* = 0 mm, Fig. 2*D*), as expected due to the cylindrical geometry.

## 2 Conclusion

Across all experiments, termites consistently consumed the core of dowels, irrespective of length or geometry. This behaviour suggests that termites exploit bite-induced echoes to locate the centre of branches and trunks, enabling them to hollow wood from the inside out. Such a strategy parallels echolocation (*17*), but in this case operates within strongly heterogeneous media, where signals are embedded in noise and function as a communication channel (*18, 19*).

This capacity may facilitate navigation within complex tree architectures while avoiding surface breaches, thereby keeping termites concealed from predators and buffered against fluctuating environmental conditions. However, the mechanisms by which termites process these signals in detail using receiving the signal through their legs and the subgenual organ, into the thoracic ganglia remain unresolved, as does the extent to which environmental noise can be filtered and what role the communication channel takes (*18*).

High-frequency waves can reduce dispersion and preserve signal integrity (*20*), and recent findings suggest that termites’ interaction with structural waves underpins their remarkable sensory capabilities (*21*), highlighting the need for investigation at higher frequencies. The cellular architecture of plant tissues may function as a natural waveguide, analogous to the bending surface waves exploited by many *Hemiptera* (*22*). However, while termites can detect direction by making use of surface waves (*13*), it remains an open question whether this is also the case for stress waves. Also, the exact localisation of sources might then be possible (*17*).

Further research on termite attraction to Acoustic Black Holes (ABHs) may uncover additional foraging behaviours and inspire novel pest-control strategies. From an applied perspective, because *Coptotermes acinaciformis* is drawn to larger timber volumes, ABHs may serve as sentinel devices near wooden structures or as lures incorporating minimal toxin, offering efficient control with reduced chemical use.

## 3 Methods

Seven colonies of *Coptotermes acinaciformis* were collected from bushland around Darwin, Northern Territory, Australia. Termite adult workers and soldiers were collected at their natural population ratio of 10:1 (*23, 24*). Our experiments included 84 setups and four repeats, and three different food-choice configurations: one short-short configuration was used as global control, one short-long configuration served twofold, to determine whether *C. acinaciformis* prefers the food resources (*3*) and as a control for the last configuration, which consisted of a combination of a long dowel and an ABH termination. In all tests, only the near identical ends of the sequentially cut wooden dowels were presented to the termites, which therefore could not measure the physical size or shape of the dowels. To visualize the cavities excavated by foraging termites, we scanned the dowels using micro-computed tomography (*25*). For the time-explicit, finite element simulations, we developed axisymmetric models of short, long, and ABH dowels using the same dimensions as in the experiment to verify the principle of wave manipulation (for details see *SI Appendix*).

## Acknowledgments

We acknowledge the technical assistance of Travers Sansom with the *μ*CT scanning of damaged dowels.

## Author contributions

S.S., J.L., and S.O. conceived this research; C.N. developed the computational models; conducted the numerical simulations; characterized the material; S.S. and S.O. designed the bioassays; S.S., T.E, and S.O. collected the specimens; S.S. conducted bioassays; C.N. and S.S. analyzed the data; all co-authors contributed to the investigation; C.N. and S.S. wrote the original draft; C.N. wrote the manuscript; C.N., J.L., T.E., and S.O. contributed to revising the manuscript; J.L., T.E., and S.O. acquired the funding; S.O. managed this project.

