## Supplemental materials for "The illusion of infinity: Acoustic black holes in wood deceive termites"

#### This PDF file includes:

Legend for Movie S1
SI References

#### Other supporting materials for this manuscript include the following:

Movie S1

### Termite collection

Seven colonies of *Coptotermes acinaciformis* were collected from the bushes around the Gunn Point road, Howard Springs, Northern Territory, 0835 Australia. Fig. S1 in (1) illustrates the staged process of termite collection during the field trip. Panels A—C depict the selection of a suitable mound, its careful dismantling, and preparation for transport. Once the internal sections containing termites were extracted, the specimens were gently knocked out of the mound material, separated from debris using sheets of carton, and placed on a runoff platform. This step was essential for isolating healthy, mature workers and soldiers—maintaining their natural population ratio approximately 10:1 using papers (2, 3), from immature instars and damaged individuals. In panel D (1), the selected termites are shown being weighed, before being placed into a PET termite jar (panel E) to establish a surrogate sub-colony. Plastic jars (85×85×130 $mm^3$) were filled with food matrix containing a mixture of water, sawdust, vermiculite and their inner own mound pieces at the ratio of about 2:1:1:4 (total of 200 g) for 30 g of healthy workers and soldiers to settle and build protective structures quickly—less than six hours for two of the colonies.

The jars were placed in a custom-built box with foam pads and tight fit compartments for transportation, then transported via air freight from Darwin to Sydney and stored in temperature and humidity-controlled laboratory cabinets (Thermoline Climatron Plant Growth CLIMATRON-360-LED-DL, 28°C, 80% relative humidity). After approximately two weeks, once the food-building matrix in the jar had been fully consumed (panel F), the material darkened to a deep brown color, indicating that the sub-colony was established and ready for use in the food choice bioassay.

### Experimental design and bioassays

Our experiments included 84 setups, one global control and two treatments for seven colonies and four repeats ($3 \times 7 \times 4 = 84$). We designed the setups based on providing access to two food resources (choice bioassay) for all 30 g of termites in one jar, through a connecting vinyl tube. T-shaped brass connector was used to give access to dowel choices for the termites coming through the tube. The first 15 mm of the dowels were supported in the brass connector and the rest of the lengths were cantilevered in the air. The experiments scheduled an initiation period of four weeks after termites were stored in the environmental cabinets in the lab, to let them eat up their food, build their own structures and get ready to forage. The termination criterion was either termites attempt to access to the outside by making a tunnel to the perimeter of the dowels or termination after a certain time period, i.e, 2 weeks (21 tests), 8 weeks (21 tests), 12 weeks (42 tests). The validity threshold was 0.3 g, due to the uncertainties in the mass measurements, including the traces of termite soil on the samples, fast increase of moisture in dry samples, and negligible termite damage to the dowels. The dowels were air-dried for 48 hours and weighed using an analytical scale (ADB 100-4, KERN & Sohn GmbH, max 120 g and increments of 0.1 mg) prior to the experiments. Upon termination, dowels were gently cleaned and air-dried for 48 hours to measure their damaged mass.

No significant difference was observed between different rounds (2-week, 8-week, and 12-week periods). Orientation of the termite jars, whether placed vertical or horizontal, made no significant difference. During the 12-week round, termites were given more foraging area by drilling a blind hole of 8mm diameter and 10 mm depth in the dowels on the cross section; however, no significant difference was observed. Except one case, in all other cases of termination due to tunneling through, we observed termites escaping/out-exploring from the top of the dowels (horizontal orientation), rather than sides or bottom. The tunnel hole was close to the supported part (brass connector rim) in 2-week and 8-week rounds, and further from the support (middle of the length) for the 12-week round. However, no significant difference was observed between these rounds with respect to the escape location.

Panel A in Fig. S2 (1) illustrates the basic configuration for treatment consisting of a dowel with an acoustic black hole (ABH) termination on one side and a solid wood dowel with a flat end on the other. The acoustic black hole is based on (4). To ensure sufficient airflow, 0.5 mm-diameter breathing holes were included at the top of the brass connector (panel B). Panel C displays all three experimental configurations: the global control (short–short), Treatment testing the food amount hypothesis (short–long), and treatment testing the reflection-reduction hypothesis (long–ABH). During the experiments, jars were placed on stainless steel trays partially filled with water, and a 30 mm thick piece of packaging foam with a larger footprint than the jars was placed underneath each one to provide insulation and dampening. The cabinet lights were turned off, and the inner glass door remained locked for the duration of the trials to ensure a stable and undisturbed environment provides an impression of the setup of the experiments in the environmental chamber.

### Wave propagation simulation

In an infinite bulk of a perfectly elastic material, a sound wave will spread out through the material as a bulk wave, decaying in amplitude because of the spread of the wavefront. However, in a finite structure such as a perfectly elastic timber dowel, the sound wave is reflected from the structure boundaries, and the energy is contained within the dowel as a guided wave, which will propagate with constant amplitude (assuming undamped, or as Movie S1 in the damped case). The complex effect of boundaries means that the energy propagates in modes that have frequency-dependent properties, which can be calculated by solution of the elastic wave equation (5). For an orthotropic material, these modes may be coupled in longitudinal and transverse directions leading to complex propagation patterns resulting from the mathematical solution to the wave equation (6). The velocity-frequency relationships of guided waves can be displayed as dispersion curves (cf. Fig. 2$C$).

Axisymmetric models of short, long and ABH dowels with the same dimensions in the experiments were developed for simulations in COMSOL Multiphysics version 6.2. Using the elastic waves time-explicit solver within Acoustics module, time-domain simulation of dowels were conducted under a pulse pressure signal centered around 2 MHz with a radially uniform amplitude of 1 Pa exerted to the end accessible to the termites. This loading condition was considered as a mechanical estimate of the elastic snap backs of the scenario with many termites biting off the wood fibers, aiming to investigate the wave propagation within the dowels. The period of the load is one eight of the elastic wave period at nominal acoustic velocity of $c = 4{,}203\ \mathrm{ms}^{-1}$ for pine cuts along the grains. The total simulation time was set as 0.3 ms to capture about six complete round-trip of the pressure wave along the length of the long dowels. We used the smeared orthotropic material model based on the measured properties of samples cut along the grains using a model updating procedure (7). In this axisymmetric model, the axial stiffness of the dowels is about four times more than that in the radial direction.

### ABH formulation for axial loading on beams

$$\rho\left(z\right) A\left(z\right) \delta_{tt} u\left(z,t\right) - \delta_z E\left(z\right) A\left(z\right) \delta_z u\left(z,t\right) = F\left(z,t\right). \qquad [1]$$

where $\rho$ and $E$ are the density and modulus of elasticity, respectively, and $A$ denotes the cross-sectional area at location $z$. This equation describes the axial displacement $u\left(z,t\right)$ as a generically nonlinear system, under a given force $F\left(z,t\right)$. Although it can be linearized under simplifying assumptions such as homogeneous material properties and geometry, we consider the nonlinearity emerging from a varying cross section $A$ in the $z$-direction to study the ABH effect. Without loss of generality, we assume the cross section is circular and the changes in the radii are given as

$$r = \gamma_r z^m \quad \text{and} \quad E = \gamma_E z^l, \qquad [2]$$

where $m$ is the exponent, denoting the ABH order, and $\gamma_r$ is a constant. Substituting Eq. 2 into Eq. 1 and assuming homogeneous material properties, i.e., $E\left(z\right) = E$, the equation of motion under free vibration, i.e., $F\left(z,t\right)$, becomes

$$\rho\left(z\right) z^{2m} \delta_{tt} u\left(z,t\right) - \delta_z \gamma_E z^{2m+l} \delta_z u\left(z,t\right) = F\left(z,t\right). \qquad [3]$$

Considering a harmonic behavior, the axial displacement can be expressed as $u\left(z,t\right) = U\left(z\right) \exp -i\omega t$ with $U\left(z\right)$ and denoting the amplitude and circular frequency, respectively, with $\omega = 2\pi f$. Assuming that variations of $E$ and $\rho$ over the length are negligible and applying spatial differentiation, Eq. 3 changes to

$$U''\left(z\right) \frac{2m+l}{\gamma_E z} U' + \frac{\omega^2 \rho}{\gamma_E z^l} = 0 \qquad [4]$$

where $U''$ and $U'$ are the first- and second-order differentiation with respect to the $z$-coordinate. Tactoring out $\omega^2$ allows the axial vibration equation with spatially varying coefficients to be solved using the Wentzel–Kramers–Brillouin approximation (8, 9). Using $\epsilon^2 = \frac{1}{\omega}$, Eq. 4 becomes

$$\epsilon^4 \left( U''\left(z\right) + \frac{2m+l}{\gamma_E z} U'\left(z\right) \right) + \frac{\rho}{\gamma_E z^l} U\left(z\right) = 0. \qquad [5]$$

As frequency increases, the $\epsilon$ tends to zero leading to a vanishing displacement response. In this case, Wentzel–Kramers–Brillouin approximation provides an asymptotically matching solution based on the singular decomposition method. One can write

$$U\left(z\right) \approx \exp \frac{i}{\delta} \sum_{n=0}^{N} S_n\left(z\right) \delta^n \quad \text{for} \quad \delta \to 0 \qquad [6]$$

where $S_n$ is the $n^{th}$ order approximation of $U$ and denotes the perturbation parameter (8, 9), and $i = -1$. By substituting Eq. 6 into Eq. 5, we transform the governing equation into a series of ordinary differential equations that can be solved in sequence, first of which gives a geometrical acoustic solution (8–10). Since Eq. 6 provides an admissible solution to Eq. 1, an acoustic black hole effect can be expected from the exponential change of thickness, cf. Eq. 2. We implemented this for a profile with $m = 2$ over a 30 mm section at the end of the ABH dowels. The dowel diameter tapers from from 19.5 mm to 1 mm, the manufacturing limit for *P. radiata* dowels.

### Damage void analysis

To visualize the voids left by termite foraging we scanned the dowels using a $\mu$CT scanner (Skyscan 2214) at a voxel resolution of 40 $\mu$m. Dowels were paired during the experiments, ensuring that each pair was exposed to termites from the same jar. This particular group was selected due to visible differences between the Long and ABH dowels .We used open-source segmentation tools (Drishti (11) and Drishti Paint (12), version 2.7) to extract the damaged regions based on grayscale thresholding. length of the dowels with the damage using a black-and-white approach. The resulting mesh was imported into Autodesk Meshmixer 3.5.474, where a solid cylinder matching the dowel's dimensions was subtracted to isolate the void volume. Final measurements were performed using the built-in tools in Meshlab.

**Movie S1. Elastic wave simulation of dowels. Pressure time-history (0 - 0.3 ms) of the 19 mm-diameter short (40 mm), long (160 mm), and ABH (160 mm) Pinus radiata dowels (modelled as bulk orthotropic material) in response to a pulse excitation (P = 1 Pa) that is uniformly distributed across the dowel end exposed to the termite feeding site.**
